# A Deep Learning Genome-Mining Strategy Improves Biosynthetic Gene Cluster Prediction

**DOI:** 10.1101/500694

**Authors:** Geoffrey D. Hannigan, David Prihoda, Andrej Palicka, Jindrich Soukup, Ondrej Klempir, Lena Rampula, Jindrich Durcak, Michael Wurst, Jakub Kotowski, Dan Chang, Rurun Wang, Grazia Piizzi, Daria J. Hazuda, Christopher H. Woelk, Danny A. Bitton

## Abstract

Natural products represent a rich reservoir of small molecule drug candidates utilized as antimicrobial drugs, anticancer therapies, and immunomodulatory agents. These molecules are microbial secondary metabolites synthesized by co-localized genes termed Biosynthetic Gene Clusters (BGCs). The increase in full microbial genomes and similar resources has led to development of BGC prediction algorithms, although their precision and ability to identify novel BGC classes could be improved. Here we present a deep learning strategy (DeepBGC) that offers more accurate BGC identification and an improved ability to extrapolate and identify novel BGC classes compared to existing tools. We supplemented this with downstream random forest classifiers that accurately predicted BGC product classes and potential chemical activity. Application of DeepBGC to bacterial genomes uncovered previously undetectable BGCs that may code for natural products with novel biologic activities. The improved accuracy and classification ability of DeepBGC represents a significant step forward for *in-silico* BGC identification.

## Introduction

Natural products are chemical compounds that are found in nature and produced by living organisms. They represent a rich reservoir of drug candidates that have proven utility across multiple therapeutic areas. Between 1981 and 2014, one third (32%) of FDA approved small molecule drugs were either unmodified natural products (6%) or natural product derivatives (26%)^1^. These include multiple classes of antibacterials, as well as oncology drugs, diabetes drugs, hypocholesterolemic drugs, and immunomodulatory agents^1,2^. The global rise in antibiotic resistance^3,4^, the increased promise of immunomodulatory agents in cancer treatment^5^, and the continued need for development of new drugs across novel and complex biology is contingent upon the identification of structurally diverse bioactive compounds^6–8^.

Early genetic work in the field of natural product discovery showed that bioactive molecules are microbial secondary metabolites whose synthesis is primarily orchestrated by genomically colocalized genes termed Biosynthetic Gene Clusters (BGCs)^9–11^. While these early insights were born out of forward genetic approaches (progressing from phenotype to sequence), the advent of next generation sequencing technologies and genomic approaches provided opportunities for reverse genetic approaches (progression from sequence to phenotype) in BGC discovery, synthesis, and characterization^2^. The surge of microbial genomic resources, including completed genome sequences of cultured and uncultured organisms, has enabled a paradigm shift in how computational methods have been used in natural product drug candidate discovery.

Numerous bioinformatics tools have leveraged the increasingly abundant genomic data to facilitate natural product genome mining^12^. Early approaches implemented simple BGC reference alignment techniques using programs like BLAST^13^, and were often paired with manual curation. Rule-based algorithms^14,15^ improved on their predecessors by using human-coded (“hard coded”) rule sets to define BGCs based on their similarity to reference genes and protein domain composition. While some recent approaches have continued to employ these “reference-based” techniques, other algorithmic advances have embraced more generalizable machine learning approaches that provide a greater ability to discover new BGC genomic elements. One such widely used machine learning approach named ClusterFinder^16^ employs a Hidden Markov Model (HMM) instead of the multiple sequence alignment based profile-HMM^17^ methods seen in other approaches such as AntiSMASH (ANTIbiotics & Secondary Metabolite Analysis SHell)^14^ and PRISM^18^.

While they have been effective, HMMs like ClusterFinder do not preserve (i.e. remember) position dependency effects between distant entities or order information^19–21^. This means HMM-based tools are unable to capture higher order information among entities^19–21^, thus limiting their ability to detect BGCs. We addressed this algorithmic limitation by implementing a deep learning approach using Recurrent Neural Networks (RNNs) and vector representations of protein family (Pfam)^22^ domains which together, unlike HMMs, are capable of intrinsically sensing short– and long-term dependency effects between adjacent and distant genomic entities^23^. This implementation yielded performance higher than another leading algorithm (ClusterFinder), including improved BGC detection accuracy from genome sequences and improved ability to identify BGCs of novel classes.

Here we introduce DeepBGC, a novel utilization of deep learning and natural language processing (NLP) strategy for improved identification of BGCs in bacterial genomes (Figure 1). DeepBGC employs a Bidirectional Long Short-Term Memory (BiLSTM) RNN^24,25^ and a word2vec-like word embedding skip-gram neural network we call pfam2vec. Compared to Clusterfinder^16^, DeepBGC improves detection of BGCs of known classes from bacterial genomes, and harnesses great potential to detect novel classes of BGCs. We supplement this with generic random forest classifiers that enables classifications of BGCs based on the product class and molecular activity of the compounds. We applied DeepBGC to bacterial reference genomes to identify BGC candidates coding for molecules with putative antibiotic activity that could not be identified using the other existing methods. In addition to bacterial reference genomes, we expect this approach to be important in microbiome metagenomic analyses, in which the improved BGC detection may empower new functional insights. To facilitate these and other analytical applications, DeepBGC is available at https://github.com/Merck/deepbgc.

**Figure 1.**
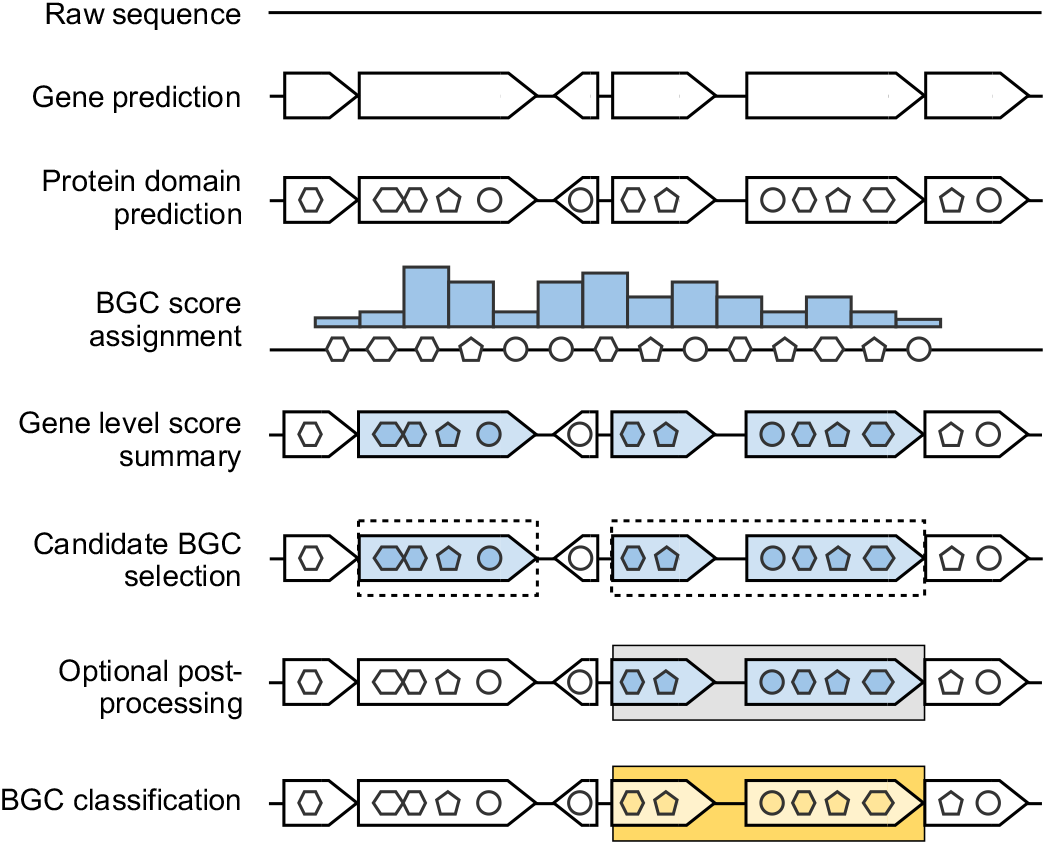
Overview of the deep learning strategy for detection of Biosynthetic Gene Clusters in bacterial genomes. (From top to bottom) raw genomic sequences (solid line) are used for gene (arrowhead structures) prediction by Prodigal ^40^. Pfam domains (circles, penta– and hexagons) are assigned to each ORF using hmmscan^17^. The BiLSTM outputs classification score (blue bars) for each domain. Domain scores are summarized across genes, which are selected accordingly (blue arrowhead structures). Consecutive candidate BGC genes are assembled to putative BGCs (dashed rectangles). An optional post-processing step allowed merging of neighboring BGC based on the presence of known biosynthetic pathway, minimum cluster length, and gaps between adjacent BGCs (gray rectangles). BGCs were classified using random forest classifiers based on compound class and molecular activity (yellow rectangles).

## Results

### Curation Yielded Diverse Training & Validation Datasets

BGC prediction is a classification task requiring the use of labelled BGC and non-BGC sequences (a “positive” and “negative” set, respectively) to train and validate the classifier. To ensure adequate comparison between the existing HMM approach and our deep learning strategy, we trained and validated our model using a training set similar to that used in Cimermancic *et al*.^16^. We built our positive training set by retrieving 617 out of 667 published labeled BGCs from Cimermancic *et al.*^16^ (Supplementary Figure S1, Table S1). We constructed a negative training set of 10128 random gene clusters based on similar principles to those described in Cimermancic *et al.*^16^. We additionally retrieved a second, supplementary dataset consisting of 1406 BGCs found in the Minimum Information about a Biosynthetic Gene cluster (MIBiG) database^26^, which were used for 10-fold cross-validation, leave-class-out validation and training random forest classifiers (Supplementary Figure S1, Table S2).

In addition to the BGC sequence datasets, we utilized whole bacterial genomes that had been manually annotated with BGC and non-BGC regions. We used the set of 9 bacterial genomes that contained 291 manually annotated BGCs from Cimermancic *et al*.^16^ and used them for validation, hyperparameter tuning, and testing (Supplementary Figure S1, Table S3). We also used a second set of 65 experimentally validated BGCs in 6 bacterial genomes (termed the validated Cimermancic *et al*.,^16^ set, Supplementary Figure S1, Table S4), which allowed for a supplemental model testing. Additionally, we retrieved a corpus of 3376 unannotated bacterial genomes that we used for generating negative samples, for pfam2vec corpus curation and for explorative application of DeepBGC to detect novel BGCs (Supplementary Figure S1, Table S5).

### Pfam2Vec Captures Biological Signal for DeepBGC Input

One major challenge in BGC identification was defining informative genomic input for the algorithm. Input sequences of biological entities can be represented at different genomic levels including nucleotides, amino acids, and genes. Of these options, sequential protein family (Pfam) domain representations have been highly informative for BGC identification^16,27^ because they represent functional elements within genes. We extended this approach by converting sequences of Pfam domain identifiers to numeric vector representations using the word2vec algorithm^28^. The resultant vectors of real numbers encapsulated domain properties based on their genomic context, allowing us to leverage contextual (and below we show functional) similarities between Pfam domains and BGCs.

We validated the ability of pfam2vec to produce functionally meaningful numeric representations of Pfam domains within genomes by calculating the average vector cosine similarity between members of superfamilies. The cosine similarities within the same Pfam superfamilies (clans) were significantly higher than the similarities between random domain vector pairs (p < 2.2 × 10^-16^, Supplementary Figure S2a). The average cosine similarity between superfamily pairs and random pairs was centered around 0 when random representation vectors were used (p = 0.09, Supplementary Figure S2b). The known domain functional annotations of the nearest domain vector pairs were more similar compared to random pairs (p < 2.2 × 10^-16^, Supplementary Figure S2c) and N– and C– terminal domain pairs appeared to be more similar in vector space compared to other domains (Supplementary Table S6). These findings suggested that pfam2vec produced functionally meaningful numeric representations of Pfam domain sequences by reflecting known superfamily similarities.

We further confirmed the functional relevance of pfam2vec vectors by evaluating their ability to discriminate BGCs by their Pfam repertoires. We accomplished this by condensing the many Pfam domains within each given BGC into a single representative BGC vector using two alternative approaches. In the first approach we created representative BGC vectors by averaging the domain vector values, and in the second approach we created a binary vector indicating the presence of each specific domain (one-hot-vector representation). We assessed the biological relevance of these two approaches by means of a t-Distributed Stochastic Neighbor Embedding (t-SNE) of all BGCs from the MIBiG database^26^, and showed that both approaches preserved similarity between BGC subclasses (Supplementary Figure S3a-b). Distinct separation of BGCs was lost when assessing their taxonomic discriminative abilities, further suggesting that BGCs were primarily defined by their functional domain architecture and not by their bacterial species (Supplementary Figure S3c-d). While one-hot-vectors meaningfully represented BGCs, they could not be used for individual domain representations, since large vectors may inflate the number of model parameters and consequently lead to model overfitting. On the contrary pfam2vec vectors provided condensed and meaningful representation of individual domains and therefore represented a functionally relevant input for RNN that could enhance BGC identification.

### Unique Model Architecture & Bootstrapping Improves BGC Prediction

The DeepBGC BiLSTM neural network was comprised of three layers: the input layer, the BiLSTM unit, and the output layer (Figure 2). The *input layer* encoded a sequence of numerical vectors representing Pfam domains in their genomic order. The *BiLSTM layer* consisted of forward and backward LSTM network layers, each consisting of a basic LSTM unit (a memory cell) with a 128-dimensional hidden state vector. The memory cell was fed with a single input vector as well as the cell’s state from the previous time step in the genome. The *output* from all LSTM memory cells was processed through a single fully connected output layer with a sigmoid activation function. This yielded a single value for each genomic Pfam entity, which represented BGC classification confidence for that Pfam domain. This model was trained using our positive (Cimermancic *et al.*^16^ labeled BGCs) and negative (random gene clusters) training sets, which were converted into their respective pfam2vec vectors. Positive and negative BGC matrices were repeatedly shuffled and concatenated to simulate real genomic context in which BGCs were scattered randomly throughout the genome and surrounded by non-BGC sequences.

**Figure 2.**
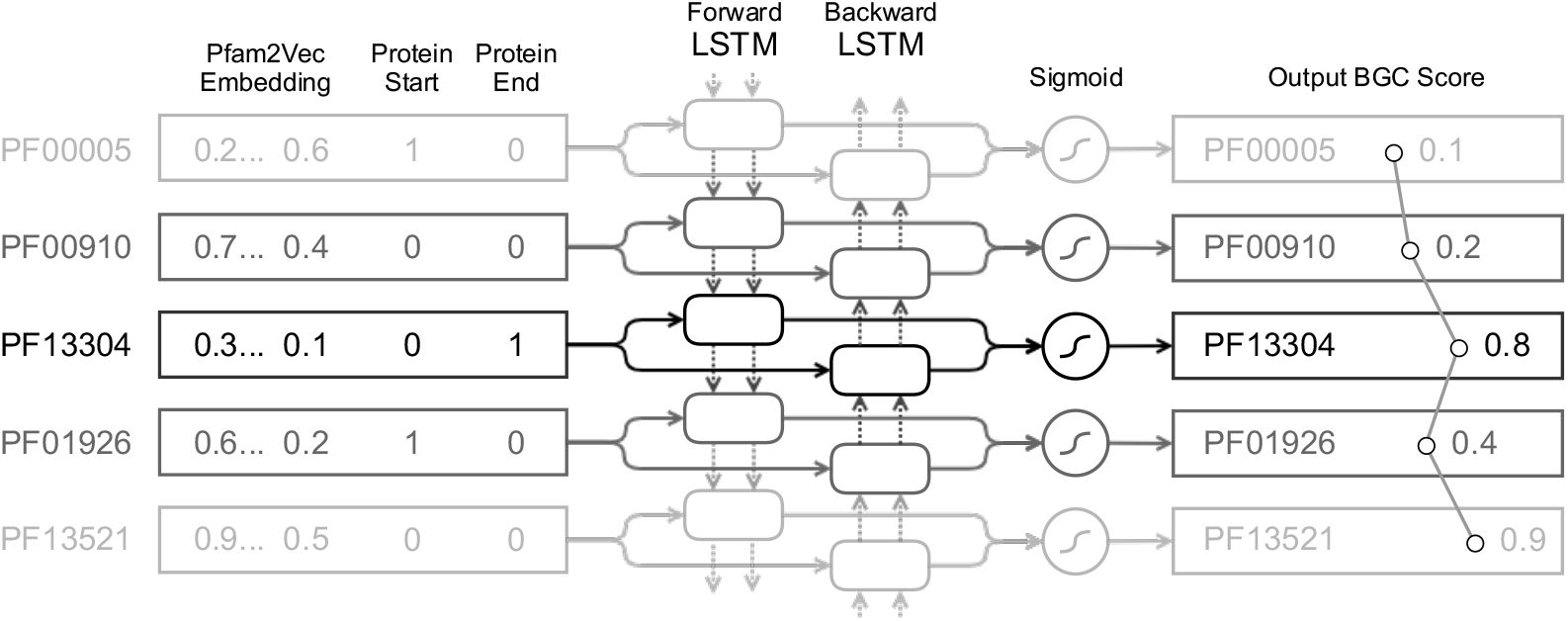
Bidirectional Long-Short Term Memory (BiLSTM) neural network architecture (left to right blocks). The network consists of three layers: input, BiLSTM network, and output layer. (top to bottom) Each row represents a time step where the BiLSTM model processes a single Pfam domain from the input sequence that is maintained in genomic order. Each Pfam domain is represented as a vector of precomputed 100-dimensional pfam2vec skip-gram embedding and two binary flags indicating whether the domain is found at the beginning or at the end of a given protein. Each LSTM memory cell receives the vector from input layer (full arrows) as well as the cell’s internal state that represents all previously seen Pfam domains (dashed arrows). The backward LSTM layer processes the vectors in reverse order, hence bi-directional. In each timestep, output from both LSTM memory cells (boxes) is processed through a single fully-connected node with sigmoid activation function (circle) that outputs a single BGC classification score for the given Pfam domain.

To compensate for the lack of a large independent validation set that could be used for optimizing model architecture, input features, and hyperparameters, we employed a bootstrap sampling technique. Because our model was trained on artificially created genomes, we bootstrapped a real-world dataset to enable hyperparameter tuning and to simultaneously prevent data leakage that would bias our estimation of accuracy in our algorithm. We performed bootstrapping using our nine manually BGC-annotated whole-genome dataset, wherein each of the 5 iterations we randomly selected 2 genomes randomly with replacement for validation and used the remaining genomes for testing (Supplementary Figure S1). We obtained an averaged Receiver Operating Characteristic (ROC) curve was obtained by combining the 5 iteration test set predictions, which revealed an improvement in precision and recall compared to the original ClusterFinder HMM model as well as a retrained version with up-to-date data (precision: 0.75, 0.28, 0.63, respectively, Figure 3a-b).

**Figure 3.**
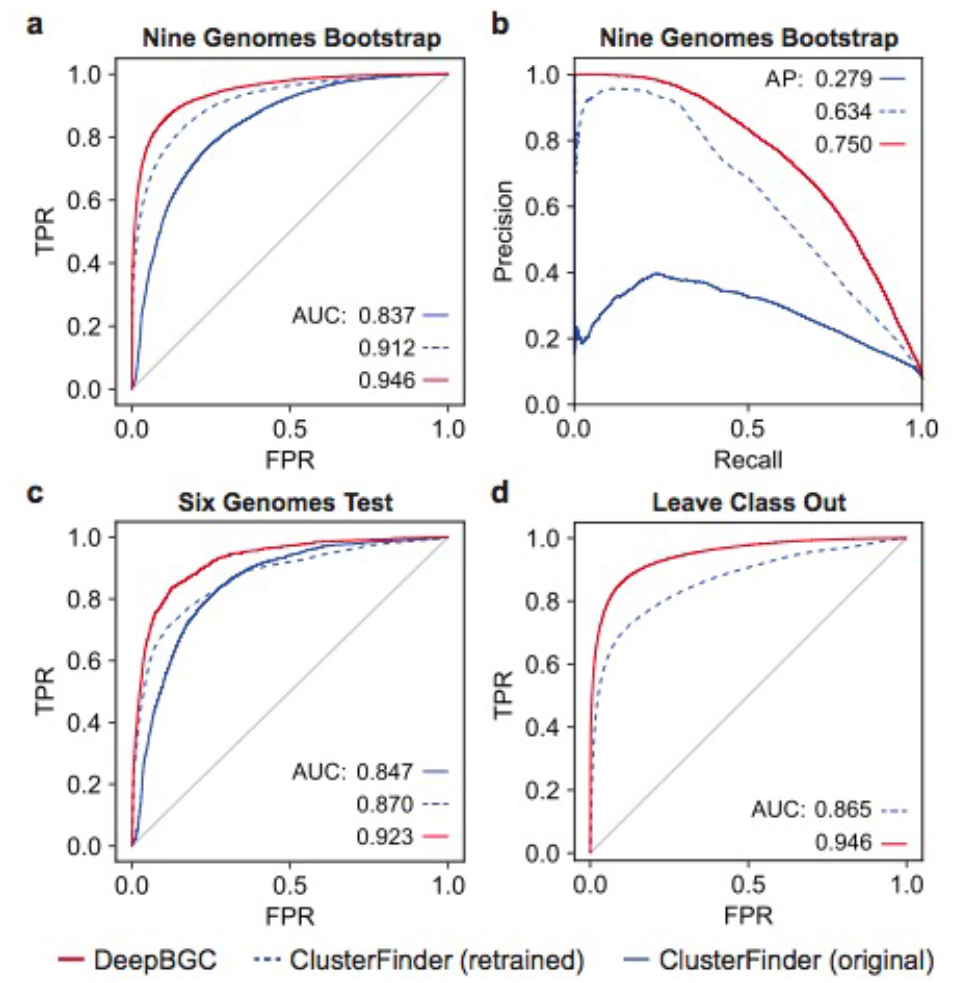
Model validation and testing on Pfam domain level using the (a) Receiver Operating Characteristic (ROC) curves and (**b**) Precision (Y-axis) Recall (X-axis) Curve reflecting performance of: (blue) original ClusterFinder HMM model, (green) ClusterFinder HMM model retrained with latest training data and latest Pfam database, and (red) DeepBGC. A total of 291 BGCs in 9 bacterial genomes were used for testing, none of them were included in the training set. The DeepBGC ROC represents combination of 5 test set predictions following bootstrap. AUC (Area Under the Curve) values are as indicated: FPR-False Positive Rate (X-axis); TPR – True positive rate (Y-axis). (**c**) ROC curves reflecting performance using a total of 65 experimentally validated BGCs that were used for testing, none of them were included in the training set. (**d**) ROC curves reflecting average performance in “Leave-Class-Out” analysis. The mean AUC for all classes is given. For individual classes performance see Supplementary Figure S5.

### DeepBGC Accuracy Outperforms Existing Models

We formally evaluated the performance of the DeepBGC model and compared it to the ClusterFinder model by testing its ability to (1) accurately identify BGC positions within whole bacterial genomes, (2) discriminate between BGCs and artificially created non-BGC sequences, and (3) identify “novel” BGC classes to which it has not been exposed. First, to address whether DeepBGC could accurately identify BGC positions within whole genomes, we evaluated the model positional accuracy using our 65 experimentally validated BGC set from six bacterial genomes^16^. This revealed an improved performance of DeepBGC (AUC = 0.923) over ClusterFinder (AUC = 0.847, Figure 3c). Second, we used 10-fold cross validation to evaluate whether our final DeepBGC model could better discriminate between BGC and artificially created non-BGC sequences compared to the existing ClusterFinder algorithm. We used BGCs in the MIBiG database^26^ as a positive set and the random gene cluster negative set of BGCs. Both sets were randomly distributed across 10 bins, with 90% of bins used for training the model (with the optimal settings) and 10% used for testing. DeepBGC (AUC = 0.984) outperformed ClusterFinder (AUC = 0.936) in differentiating between positive and negative samples (Supplementary Figure S4). Third, to evaluate DeepBGC’s ability to identify “novel” BGCs, we carried out a “Leave-class-out” validation in which we assessed the models’ abilities to identify a single BGC class in the test set that was intentionally omitted from the training set. DeepBGC yielded more accurate identification of classes it had not encountered (AUC = 0.946) compared to ClusterFinder (AUC = 0.865; Figure 3d, Supplementary Figure S5). Overall DeepBGC yielded improved BGC identification accuracy, and was better able to accurately extrapolate to identify BGCs of classes it had not encountered previously.

### DeepBGC Yields Improved Precision & BGC Coverage

The most common use case for BGC identification models such as DeepBGC is locating BGCs within bacterial genomes. It was therefore important for us to validate DeepBGC as having improved positional predictive accuracy of BGCs, in addition to an improved precision and recall.

We achieved this by comparing the accuracy of DeepBGC and ClusterFinder BGC detection in a subset of our manually annotated bacterial genomes. To account for potential differential impacts of false positive and true positive rate thresholds, we applied two distinct cutoffs based on the respective domain level ROCs. We accomplished this by applying a stringent domain level False Positive Rate (FPR) of 10% as well as a lenient cutoff of 80% True Positive Rate (TPR, Supplementary Table S7, Supplementary Figure S6a-b).

With the stringent cutoff of 10% FPR, the number of BGCs predicted by DeepBGC was consistently higher than those predicted by ClusterFinder, regardless of the BGC coverage threshold (Figure 4a-b). ClusterFinder displayed a sharp decline in the number of predicted BGCs as the coverage threshold increased (Figure 4b), indicating that BGCs predicted by this approach were typically short (Supplementary Figure S7). The overall precision at the BGC level remained comparable between the two models (precision = 0.34, 0.26, Figure 4c). Under a more lenient TPR threshold of 80%, ClusterFinder predicted more BGCs than DeepBGC when coverage threshold remained less than 68% (Figure 4d-e). But we found that ClusterFinder predictions were of low precision and were composed of many false positives (Figure 4f, BGC level precision < 0. 09, Supplementary Figure S8). On the contrary, DeepBGC displayed > 4-fold increase in precision compared to ClusterFinder (precision > 0.4, Figure 4f).

**Figure 4.**
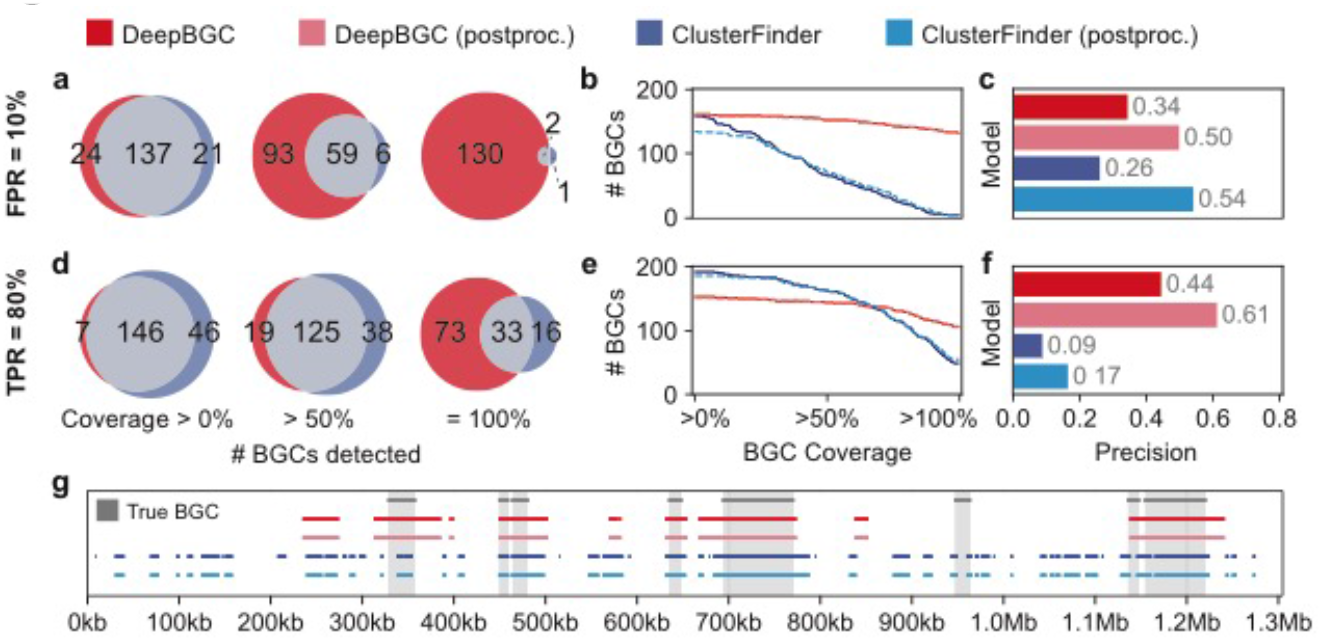
Precision and coverage of DeepBGC and ClusterFinder algorithms. (**a**) Number of true BGCs detected by DeepBGC (red), ClusterFinder (blue) and both models (grey), based on three BGC coverage thresholds: any (> 0%), majority (> 50%), and full (100%). Coverage of each annotated true BGC is defined as the fraction of its nucleotide sequence overlapping with colocated predicted BGCs. The first bootstrap test split of 7 out of 9 genomes was used for comparison. Domains were retrieved based on a fixed False Positive Rate (FPR) of 10%. Genes containing candidate Pfam domains were summarized to produce putative BGCs that were compared to the actual BGCs in the split data. (**b**) Cumulative coverage plot of actual BGCs by predicted BGCs for DeepBGC (red) and ClusterFinder (blue) also following post-processing (dashed). (**c**) BGC level precision for DeepBGC (red) and ClusterFinder (blue) also following post-processing (light colors) at FPR 10%. Precision was calculated as follows: the number of true positives (any overlap between actual and predicted BGCs) divided by total number of predicted BGCs. (**d-f**) Same as ‘**a-c**’ but at 80% TPR cutoff (**g**) A snapshot of contig view (X-axis genomic coordinates of *Micromonospora sp*.), true predictions (grey shade and bar), ClusterFinder raw and post-processed predictions (dark and light blue), DeepBGC raw and post-processed (dark and light red). For simplicity only part of the contig is shown and only at 80% TPR threshold. For all contigs, thresholds and models, see Supplementary Figure S7-8.

To correct for short BGCs predicted by ClusterFinder, Cimermancic et al.^16^ applied a postprocessing step whereby neighboring clusters were merged if they were separated only by a single gene. Putative clusters that were below 2Kb in length, as well as those not containing a known biosynthetic domain, were filtered out. We implemented this approach and found that while this step dramatically improved precision for both models (Figure 4c,f), it also inevitably removed a subset of true positive predictions, most notably for ClusterFinder at 10% FPR (Figure 4b). We thus concluded that DeepBGC not only reduced the number of false predictions compared to ClusterFinder, but it also located BGCs within genomes more accurately.

### Random Forests Provide Product & Activity Classification

To identify the biosynthetic products derived from predicted BGCs, we classified BGC sequences by training and testing a random forest classifier using the MIBiG database, which contains classification of BGCs to one or more compound classes (Table 1). Nested five-fold crossvalidation revealed that our random forest classifier exhibited an average AUC of 0.80, broadly comparable with the BGC classifier implemented in antiSMASH (AUC = 0.78, Table 1, Supplementary Figure S9). Our approach can also reveal the most influential Pfam domains that drive the classifier decisions (Supplementary Figure S10). Our DeepBGC random forest classifier therefore provided a data-driven alternative to the rule-based antiSMASH classification approach.

**Table 1.**
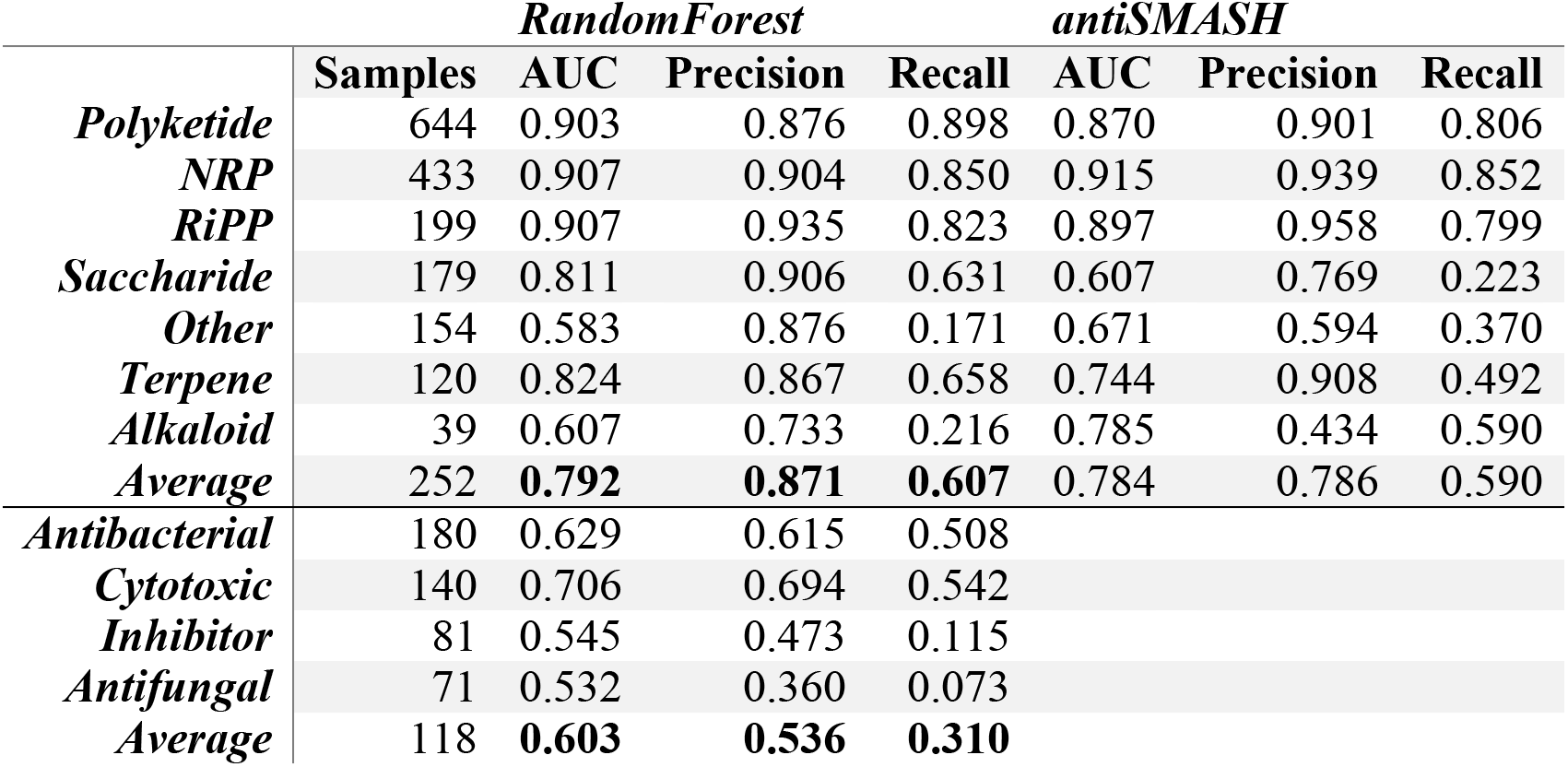
Random forest classifiers and antiSMASH performance for classifying BGCs based on their products and their underlying activity. The classifier was trained using 1355 MIBiG labeled BGCs belonging to one or more compound classes including Polyketides (PKS), Non-Ribosomally synthesized Peptides (NRP), Ribosomally Synthesized and Post-translationally modified Peptides (RiPP), Saccharides, Terpenes, Alkaloids, and those belonging to other rarer classes (“other”). Areas Under the Curve (AUC) was determined using 5-fold cross-validation. Respective confusion matrix and important domain features are provided in Supplementary Figure S9-S10. For molecular activity classification random forest was used as before on 370 molecular activity labeled MIBiG BGCs. Only antibacterial, cytotoxic, inhibitor or antifungal classes are accounted for.

In addition to identifying BGC classes, we also evaluated our ability to predict BGC molecular activity information using the 370 molecular activity labeled MIBiG BGC subset. Due to the small sample size, the classifier accounted only for the four most common compound activity classes: antibacterial, cytotoxic, inhibitor, and antifungal. Using 5-fold cross-validation, BGCs were classified according to their compound activity with modest precision (average AUC 0.61, Table 1). Larger training sets will be needed for improved performance in future work.

### DeepBGC Predicts Novel Antibacterial BGCs

Above we showed that DeepBGC, together with random forest classifiers, could effectively identify BGCs and classify their compound class and molecular activity. We therefore applied this model to unearth novel BGCs that could not be predicted by other approaches. We accomplished this by using a bacterial reference set of 3376 RefSeq bacterial genomes^29^ and subjected these to DeepBGC, antiSMASH and ClusterFinder analyses, followed by a systematic comparison of their predictions. To avoid an artificially inflated number of putative novel predictions, we maximized the ability of ClusterFinder and antiSMASH to detect BGCs by accepting their default (lenient) settings, while conversely applying a strict cutoff only for DeepBGC (2% FPR at the domain level). Under these criteria, ClusterFinder predicted > 4.5 times more BGCs (62491) than antiSMASH (13865) and > 5.5 times more than DeepBGC (10926). As expected, the majority of BGCs that were identified by ClusterFinder (~75%) could not be identified by DeepBGC or antiSMASH (Figure 5a). ClusterFinder predictions showed comparable overlap with DeepBGC and antiSMASH (18% and 15% respectively). On the contrary, DeepBGC comparison revealed that the majority of DeepBGC predictions overlapped with ClusterFinder (~55%), and ~39% with antiSMASH of which 35% overlapped with both (Figure 5a). Only ~5% (566) of DeepBGC predictions could not be uncovered by any other method. Interrogating the respective class and species distributions of these putative novel BGCs alongside all DeepBGC predictions revealed that DeepBGC clusters were enriched for BGCs with no confident class (~70%) and for those that originated from the *Mycobacterium* genus (~49%, Supplementary Figure S11).

We further explored these novel signatures by interrogating the ~5% novel BGCs that DeepBGC detected. We accomplished this by evaluating 227 BGCs that consisted of at least 5 Pfam domains and classified them based on their compound class and molecular activity. The novel BGCs that could not be confidently assigned to a single compound class straddled the borders between distinct classes (grey plus signs, Figure 5b), while the remaining were tightly clustered with known BGCs according to their respective class (Figure 5b).

**Figure 5.**
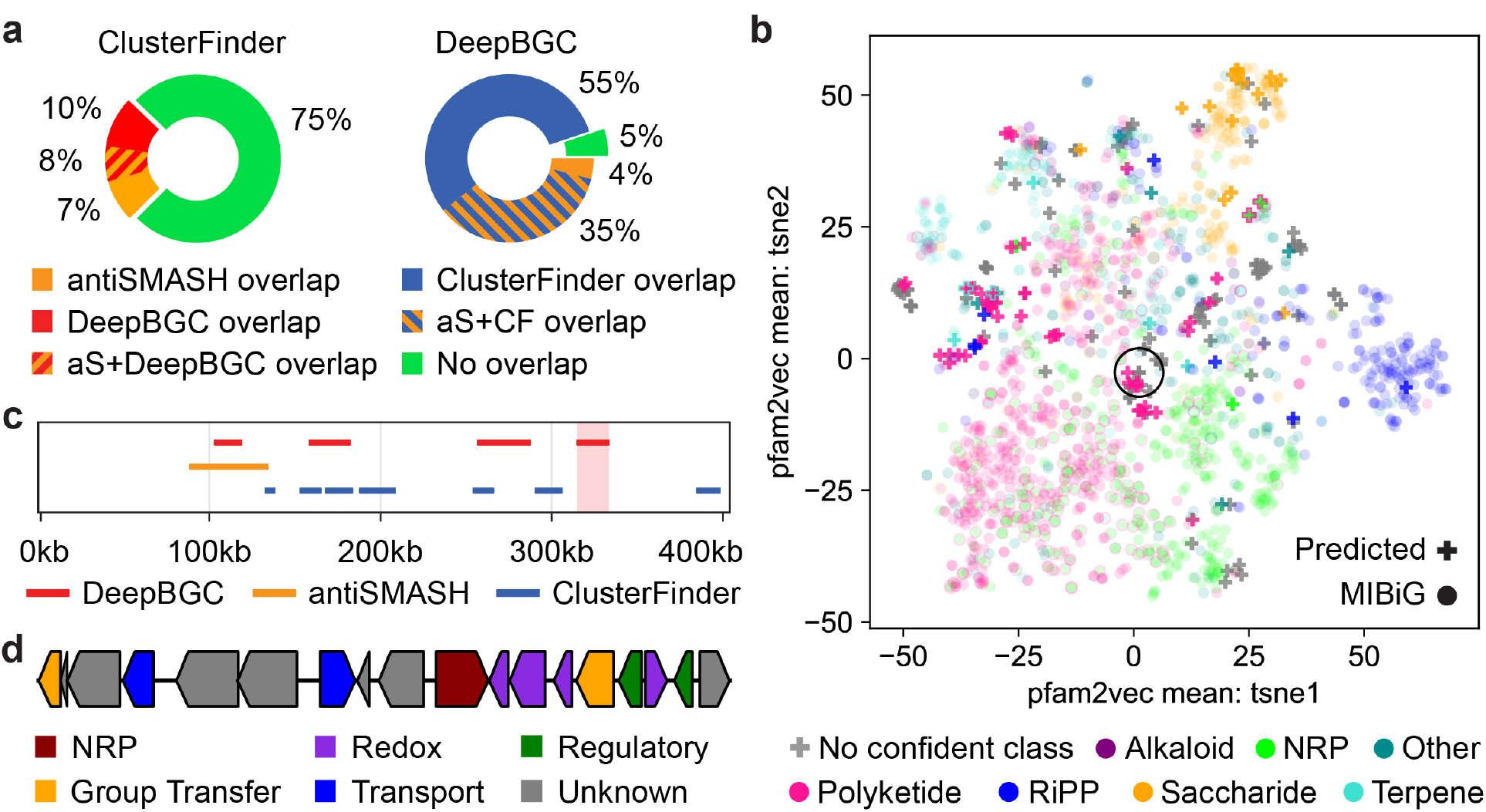
DeepBGC uncovers novel BGCs with antibacterial activity in bacterial genomes. (**a**) Comparison of BGC predictions between (left) ClusterFinder and (right) DeepBGC. Default ClusterFinder settings from antiSMASH suite were used. In DeepBGC, a 2% FPR at the domain level was applied with no further post-processing. Overlap of predictions of each model with antiSMASH rule-based predictions (default settings) is also given. (**b**) t-Distributed Stochastic Neighbor Embedding (t-SNE) of all 1355 class labelled BGCs from the MIBiG database (circles) overlaid with the putative novel 227 BGCs that could be predicted solely by DeepBGC (plus signs). BGCs were represented by the mean value of their pfam2vec domain vectors and are colored by the respective known or predicted class as indicated. (**c**) A snapshot of contig view (X-axis genomic coordinates of *Mycobacterium tuberculosis*) of BGC predictions by (red) DeepBGC, (blue) ClusterFinder, and (orange) antiSMASH combined with ClusterFinder. A novel BGC candidate predicted only by BGC is highlighted (light red shade). (**d**) The novel BGC structure is given, respective genes are colored based on the underlying domain type. For domain IDs see Supplementary Table S9.

To begin evaluating the individual BGCs, we ranked our novel predictions based on their total number of Pfam domain counts, their predicted antibacterial activity score and their similarity to known BGC in the MIBiG database (Supplementary Table S8). We used this list to manually identify a candidate BGC that could not be assigned to a specific compound class, displayed a low similarity score to known BGCs and had a high antibacterial prediction score. This novel BGC resided in the genome of the pathogenic bacterium *Mycobacterium tuberculosis*, in which BGCs are known to be abundant^30–34^. The cluster was distant from any other neighboring predictions (~9kb Figure 5c) and rich in a diverse set of regulatory, transport and modifying enzymes (Figure 5d, Supplementary Table S9) including acetyltransferase and glyoxalase/bleomycin resistance protein, which have been known to catalyze diverse biochemical reactions^35^. The cluster also encoded for a type II toxin-antitoxin system, further supporting its potential cytotoxic activity^36^. Such wealth of modifying enzymes could potentially grant the final natural product a novel chemistry. A search for BGCs with similar domain architecture revealed a similar cluster (80% similarity) in a different *Mycobacterium tuberculosis* strain. More divergent clusters with >60% similarity scores were discovered in other *Mycobacterium* species, and were also supported by predictions from antiSMASH and ClusterFinder (Supplementary Figure S12). Together these discoveries highlight the value of DeepBGC and its ability to mine bacterial genomes to provide previously unrealized insights into bacterial natural product chemistry.

## Discussion

Here we present DeepBGC, a comprehensive deep learning strategy for identifying BGCs from bacterial genomes and classifying them by their product class and chemical activity. Our deep learning approach is based on concepts from the NLP field and builds on existing algorithms that either suffer from a restricted ability to identify novel BGC classes or from a limited ability to accurately identify BGCs within a genome. We demonstrated that our DeepBGC approach outperformed one commonly used machine learning algorithm, ClusterFinder, in its ability to identify BGCs accurately within a genome (Supplementary Table S10). Our leave-class-out analysis suggested that the DeepBGC model also possessed a greater potential to extrapolate and identify BGC classes that it has not encountered before. The supplemental random forest classification approach allowed us to accurately identify BGC classes by their protein family (Pfam) domain composition, and enabled some prediction of the chemical activity of the resulting secondary metabolites despite the limited sample size available for training. Finally, like other machine learning algorithms, DeepBGC is poised to continue improving over time with the continuous discovery, validation, and labelling of BGCs in microbial genomes.

Machine learning has had a dramatic impact on NLP methodologies, giving rise to powerful word embedding techniques such as word2vec, which allow representation of words as low-dimensionality vectors of real numbers through which they enhance learning by context^37^. Asgari *et al.*^38^ recently adopted word embedding and neural networks to improve classification of protein families using amino acid sequence vectors and Kim *et al*.^39^ has also introduced Mut2Vec for representation of cancerous mutations. In our current study, we fortified our DeepBGC BiLSTM network’s ability to learn complex patterns in genomic sequences with a novel pfam2vec representation vectors that were generated from a large unlabeled corpus of genomic sequences. By doing so we enabled an improved machine understanding of the enigmatic genomic context. We believe that our pfam2vec approach could further assist in annotating domains of unknown functions based on their genomic context. To this end, we also provided a set of nearest domain pairs of known and unknown functions (Supplementary Table S11), yet this pursuit was ultimately beyond the scope of this study and we will continue pursuing this in future work.

We applied our model to a real-world dataset (collection of reference bacterial genomes) to highlight its ability to provide unique insights into bacterial BGCs. The inflated number of BGCs predicted by ClusterFinder could be readily explained by its low precision. Our results suggested that many ClusterFinder predictions were false positives, and those that were not false positives only represented small fractions of true BGCs. Throughout the study, parameter settings were conservatively chosen because we preferred underprediction over annotation of bacterial genomes with incorrectly predicted BGCs.

We found that our model identified BGCs of diverse product classes that ClusterFinder could not identify, although the majority of BGC classes could not be confidently assigned. This suggests a potential for identifying novel BGCs, warranting significant future validation to explore those BGC candidates. These unknown BGCs were largely found within Mycobacteria, a genus known to harbor many diverse BGCs ^30–34^ Therefore, our model not only identifies potential new BGC signatures, but it does so in bacteria with a known prominence for BGCs, and thus the bacteria we might expect a priori to have a significant reservoir of novel BGCs.

While we illustrated the application of this tool to reference genomes, we also anticipate DeepBGC applications to the microbiome through shotgun metagenomic datasets. An understanding of differential BGC presence or expression (using meta-transcriptomics approaches) could provide new insights into microbiome functionality, underlying mechanisms of disease, and therapeutic approaches. Although beyond the scope of this work, the incorporation of DeepBGC into microbiome sequence analyses is an exciting avenue for future studies.

While our deep learning based, DeepBGC approach outperformed other commonly used models, it is important to note its limitations. Like other existing models, this model was trained on existing BGC databases that are heavily biased towards BGCs from natural product “workhorses” such as *Streptomyces*. This bias in the training data is likely to limit the ability of the model to identify novel BGCs in bacterial sources that are poorly characterized in the databases, including bacteria found in complex microbial communities (the microbiome). We addressed this extrapolation concern by performing leave-class-out validation to highlight the generalizability of our approach over other existing approaches. Despite our improved performance, further work is needed to curate more diverse BGC databases which can be used to improve the training and validation, and overall performance as a result, of our model.

Despite this limitation, our model represents a significant step forward for natural product discovery. Due to our model’s improved ability to identify novel BGCs, we showed that we could identify BGC that were missed by other existing models, and thereby identify previously unknown sources for natural products in existing bacterial genome sequences. By reducing the number of fragmented BGCs being identified in bacterial genomes, our improved prediction accuracy will reduce BGC count inflation. Additionally, by providing more accurate BGC border predictions, we will reduce the human triage effort of cleaning up predicted BGCs whose genomic positions were not entirely accurate. Together DeepBGC represents an advancement over the current “state-of-the-art” by improving BGC identification accuracy, BGC genomic location prediction, and identification of potentially novel BGC signatures that were not present in the current training knowledgebase. This will be used to empower follow up genome mining for novel BGCs and their resulting natural products, and the improved extrapolation capabilities will empower BGC mining of microbiome datasets, which still represent an under-explored genomic BGC resource. These microbiome analyses, including associations with disease phenotypes and identification of novel chemical matter in classes such as antibiotics or immunomodulatory agents, could have important clinical impacts for translating microbiome data to therapeutic interventions.

## Online Methods

### Data & Code Availability

All data used in this work was obtained from the public domain and is specified in the respective methods sections. All code for this publication is available at the following GitHub repository: https://github.com/Merck/bgc-pipeline. DeepBGC is available for installation and use as a Python package at the following GitHub repository: https://github.com/Merck/deepbgc.

### Open Reading Frame Identification

Open reading frames were predicted in 3376 reference bacterial genomes^29^ using Prodigal^40^ version 2.6.3 with default parameters. All other sequences were downloaded with annotations and gene locations.

### Protein Family Identification

Protein family domains were identified using HMMER^17^, HMMScan version 3.1b2, and the Pfam database version 31^22^. This Pfam database was used for all applications except for the original ClusterFinder algorithm, where it was preserved at legacy version 27. Hmmscan tabular output was filtered using BioPython SearchIO module (version 1.70) to preserve only highest scoring Pfam regions with e-value < 0.01. The resulting list of Pfam domains was sorted by the gene and the domain start location.

### Pfam2vec Implementation

Pfam2vec embedding was generated using the original word2vec implementation^28^ wrapped in the word2vec python module (version 0.9.2). After bootstrap evaluation, the following hyperparameters were chosen: 100 dimensions, 8 training iterations and skipgram architecture. The training corpus consisted of 3376 documents and 23425967 words (15686 unique Pfam identifiers). Each document contained a space-separated list of Pfam identifiers representing all Pfam domains of a specific bacterial genome maintained in their genomic order. To evaluate the pfam2vec embedding, cosine similarity of domain vectors were used in comparison with domain membership to one of 604 Pfam superfamilies (clans) from the Pfam database version 31^22^. First, cosine similarity of all (non-identity) pairs of domains from all Pfam superfamilies was compared to cosine similarity of all pairs from Pfam superfamilies that were randomly shuffled (Wilcoxon rank-sum test). Second, the same calculation was performed with pfam2vec vectors replaced by random numeric vectors (Wilcoxon rank-sum test). Third, Levenshtein distance of Pfam domain descriptions was calculated for each pfam2vec vector and its nearest neighbor by cosine similarity and compared to the description similarity of each pfam2vec vector and a randomly selected pfam2vec vector (Wilcoxon rank-sum test). Finally, average pfam2vec vector representation of each BGC from the MIBiG set was calculated as an average of its list of pfam2vec vectors, with averages computed separately for each dimension. The vector representation was reduced to two dimensions for visualization using the scikit-learn manifold.tSNE package with cosine metric, random initialization and default perplexity of 30.

### DeepBGC Training Set

DeepBGC was trained on a subset of the original ClusterFinder positive training set and on a negative set generated based on similar principles as the ClusterFinder negative training set. To collect the original positive training set, 681 accession IDs were obtained from the ClusterFinder supplementary table and searched on NCBI, which returned 617 sequences. To generate the negative set, we collected and preprocessed a public reference set of 3376 bacteria. For each reference bacterium, regions similar to known MIBiG BGC (version 1.3) were removed (Blastn^13^ with 95% threshold). To generate a single negative sample, a random reference bacterium and a random sample from the positive ClusterFinder set were selected. Each gene in the positive sample was replaced with a random gene from the reference bacteria, while considering only 1% of genes that were most similar in number of Pfam domains. In total, 3 samples were generated from each reference bacteria, producing 10128 negative samples.

### DeepBGC Implementation

The BiLSTM model was implemented using Keras (version 2.1.6) with TensorFlow backend (version 1.6.0). The architecture consisted of a single Keras Sequential model with two layers. First, the model contained a stateful BiLSTM layer with 128 units and dropout of 0.2. Second, the model contained a time-distributed Dense layer with sigmoid activation and 1 output unit. The input was a sequence of Pfam domains represented by 102-dimensional vectors consisting of the 100-dimensional pfam2vec embedding and two binary flags marking domains found at the beginning or end of proteins. The output was a sequence of values between 0 and 1 representing the prediction score of given domain to be part of a BGC. In each training epoch, all positive and negative samples were shuffled randomly and concatenated to create an artificial genome. Training was configured with 256 timesteps and a batch size of 64. Thus, the training sequence of each epoch was separated into 64 subsequences, each trained in parallel in batches of 256 timesteps, processing a single training vector at each timestep. The final model was trained for 328 epochs using the Adam optimizer with learning rate of 1e-4 and weighted binary cross-entropy loss function (weights are inversely proportional to size of positive and negative samples in our training dataset).

To obtain BGC regions used for BGC-level analysis, first, predictions were averaged in each gene, BGC genes were selected using any given threshold and consecutive BGC genes were merged. Optionally, postprocessed BGC regions were created by applying filters defined in Cimermancic *et al.*^16^ : merging BGC regions at most one gene apart and filtering out regions with less than 2000 nucleotides and regions with no known biosynthetic domains from the current list of 133 domains published in the ClusterFinder^16^ submodule of antiSMASH^14^.

### DeepBGC Validation

The primary evaluation metric published in Cimermancic *et al*.,^16^ was a ROC curve based on 10 reference genomes that are fully annotated with BGC and non-BGC regions. The associated genomes were retrieved based on the list of gene loci provided in the supplementary table^16^. By querying the gene loci on NCBI, 9 out of 10 of the original genomes were obtained. The original *Streptomyces roseosporus* genome could not be retrieved. In three of the genomes the genes were not found in a single contig, but in multiple contigs (2 for *Streptomyces ghanaensis*, 2 for *Streptomyces sp. AA4* and 3 for *Streptomyces sp. C*). The sequences were updated since the release of the paper, which resulted in a location shift of 7% of genes and removal of 10 BGC genes from the *Streptomyces pristinaespiralis* genome.

For DeepBGC training we use positive and negative samples described above. Since this dataset is artificially created it lacks some features of real-world data (potential nonrandom distribution of BGC across the genome, real distribution and order of the genes etc.). Therefore, there was no guarantee that model trained on our artificial training set will perform with the same accuracy on real genomes. To address that we chose to tune hyperparameters using a subset of real world data. To avoid the reduction of our validation set and prevent a leakage from training set to validation set we chose a ‘bootstrap approach’. It consists of creating multiple models, where each model will utilize a small part of validation set for hyperparameter tuning and the rest for testing. Averaging the results of those multiple models’ results is proven to converge to the unbiased estimation of the accuracy.

During hyperparameter tuning we considered following parameters and values: learning rate (0.001, 0.0001), number of pfam2vec training iterations (4, 8, 16, 32), number of pfam2vec dimensions (50, 100, 200) and positive training sample weight (1, 16.415). Keras EarlyStopping method was used with minimum delta of 0.0005 on validation ROC AUC within 100 epochs. We realized that majority of the learned models, regardless of the validation genome selection, preferred following parameters: 0.0001 for learning rate, 8 pfam2vec training iterations, 100 pfam2vec dimensions and positive sample weight based on the negative/positive ratio of 16.415.

BGC-level coverage evaluation was performed on the test set of the first bootstrap split. Predictions of each model were converted into BGC regions (with and without postprocessing) using the method defined above. True BGC coverage of each model was calculated for each annotated BGC region as the fraction of the true BGC region covered by all its overlapping predicted BGC regions of given model. A BGC was considered detected when its coverage was above any given threshold. Coverage distribution was calculated by evaluating all coverage thresholds from 0% up to 100% in steps of 0.1%. Next, each predicted BGC region is marked as true positive if it overlaps with a true BGC region and as a false positive if it does not. Finally, BGC-level precision was calculated as the number of true positive regions divided by the total number of predicted regions.

The secondary evaluation metric published in Cimermancic *et al*.^16^ was a True Positive Rate (TPR) evaluation based on 65 BGCs in their genomic context of 6 genomes. All BGC locations were provided along with names of source bacteria in the supplementary table in Cimermancic *et al*.,^16^. First, genomes were found on NCBI by manually querying the organism names. Second, BGC start and end locations were validated to match with start and end locations of annotated genes present in the retrieved genomes. Finally, we obtained BGC predictions using DeepBGC, original ClusterFinder and retrained ClusterFinder. The original evaluation metric was based on calculating TPR in terms of fraction of BGCs detected with median ClusterFinder HMM prediction above 0.4 threshold. After the inspection of ClusterFinder predictions it was found that more than 42% of the domains outside the defined BGC regions were detected above the given threshold, relying on heavy further postprocessing and manual annotation to filter out false positives. Therefore, the sequences were evaluated using a ROC curve, which considers all unannotated regions to be negative, producing a lower bound of the AUC value which is unbiased to either of the two models.

To perform cross validation and leave-class-out validation we obtained all 1406 BGC samples from MIBiG (version 1.3) and our negative set of 10128 samples. Each sample was represented as a sequence of Pfam domain identifiers. For cross validation, samples were randomly distributed into 10 splits. In each of the 10 cross-validation folds, models were trained on 9 splits and evaluated on 1 split. Training and testing samples were shuffled and concatenated to create artificial genomes. An average ROC was computed by concatenating all test split predictions. In leave-class-out validation, 1003 samples from 6 non-hybrid classes (Polyketide, NRP, RiPP, Saccharide, Terpene, Alkaloid) were selected. For each class, the models were trained on all other classes and random two thirds of negative samples. Thereafter, models were tested on a given class (upsampled to 500 samples by sampling with replacement) and the remaining third of negative samples. Again, training and testing samples were shuffled and concatenated to create artificial genomes. This was performed three times for each class with different random splits and random shuffles to minimize the influence of any random initialization. An average ROC was computed by concatenating all test predictions.

### ClusterFinder Implementation

ClusterFinder predictions were produced using antiSMASH (version 4.1.0) with ClusterFinder enabled and with default parameters. Raw prediction scores for each Pfam domain were parsed from the final Genbank output files. These scores were used to produce domain-level ROC curves. Raw and postprocessed BGC regions used for BGC-level analysis were obtained as described earlier.

To retrain the model for cross validation and leave-class-out validation, the ClusterFinder HMM was reimplemented using the hmmlearn python module (version 0.2.0). The transition and starting probability matrices of the original model were used. The emission probability matrix was recomputed using the new positive and negative training set preprocessed with Pfam database version 31.

### Random Forest Multi-Label Product Classification

Biosynthetic product class and activity training data were obtained from the MIBiG database (version 1.3) in JSON format. Product classes were extracted from the “biosyn_class” field, producing 1355 labelled training samples. Product activities were extracted from the “chem_act” field of each compound in the “compounds” field, excluding BGCs with no known product activities, producing 370 training samples. A separate random forest classifier was trained for both domains using the scikit-learn python module (version 0.19.1). The classes were predicted using multi-label classification, where each sample is labelled with a binary vector representing presence of zero or more classes. Global feature importance was obtained using native scikit-learn method, class-specific feature importance was calculated by training a separate temporary random forest classifier for each class. To evaluate model performance, 5-fold cross validation was used, producing 5 sets of real-valued prediction scores which were merged and compared with expected output at different thresholds to produce a ROC curve. A confusion matrix was generated by treating each occurring combination of biosynthetic classes as a single separate hybrid class. To produce the antiSMASH ROC curve, all MIBiG BGC genbank files were processed through antiSMASH with default parameters. The resulting BGC classes were parsed from the text output files and converted to the output binary vector, which was used to generate a ROC curve with a single threshold.

## Acknowledgments

We thank Otakar Smrz, Joseph Lehar and Ivo Lasek for stimulating discussions. We are immensely grateful to David Dzamba, Matthew Tudor, Jyoti Shah, Petr Mejzlik, Jakub Arnold and Gergely Temesi for their comments on earlier version of the manuscript. We are also especially grateful to Mohamed Donia for the stimulating discussions and comments. We also would like to acknowledge and thank Nicole L. Glazer, Jens Christensen, and Carol A. Rohl for supporting this work.

## Author Contributions

GH, CW and DB conceived, designed and supervised the study. JS and LR guided the implementation of BiLSTM network by AP. JD, JS and DP explored protein and BGC similarities approaches. JS and DB designed the pfam2vec algorithm, JS implemented it and DP evaluated its performance. JS designed the bootstrap approach and DP implemented it. JK, MW, and DC advised and assessed the HMM, LSTM and classification methods. DP developed and implemented the entire pipeline and performed all data analyses in this study. DP, OK and JS designed and implemented the random forest classifier. RW evaluated DeepBGC predictions and GP and DJH advised throughout the study. DP and DB designed the figures. DB drafted the manuscript. GH, CW, RW, and DP contributed to the final draft, all authors read and approved the final version of the manuscript.

