## Supplementary Figures for "A Deep Learning Genome-Mining Strategy Improves Biosynthetic Gene Cluster Prediction"

### Supplementary Material

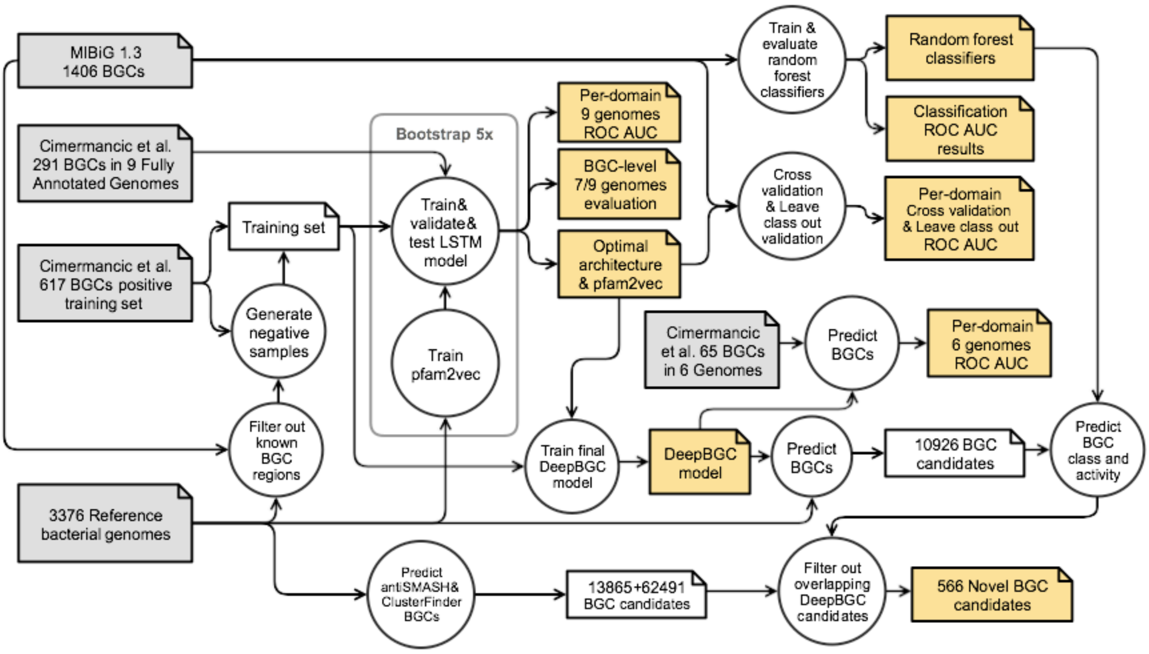


**Supplementary Figure S1.** Diagram of the major steps in DeepBGC analysis, including the relevant datasets used and the progression of algorithms and their performance tests. Input datasets are shown in grey, published results are shown in yellow.

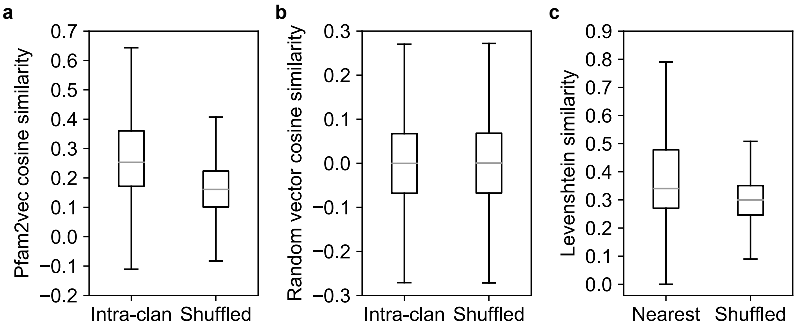


**Supplementary Figure S2.** Pfam2vec captures similarity between domains. (**a**) Cosine similarity between pairs of domains within the same clan (intra-clan) and between random pairs (shuffled) (**b**) Cosine similarity between intra-clan and shuffled pairs using random numeric vectors (c) Levenshtein distance of domain descriptions of nearest known neighbor domain by pfam2vec cosine similarity versus a random domain.

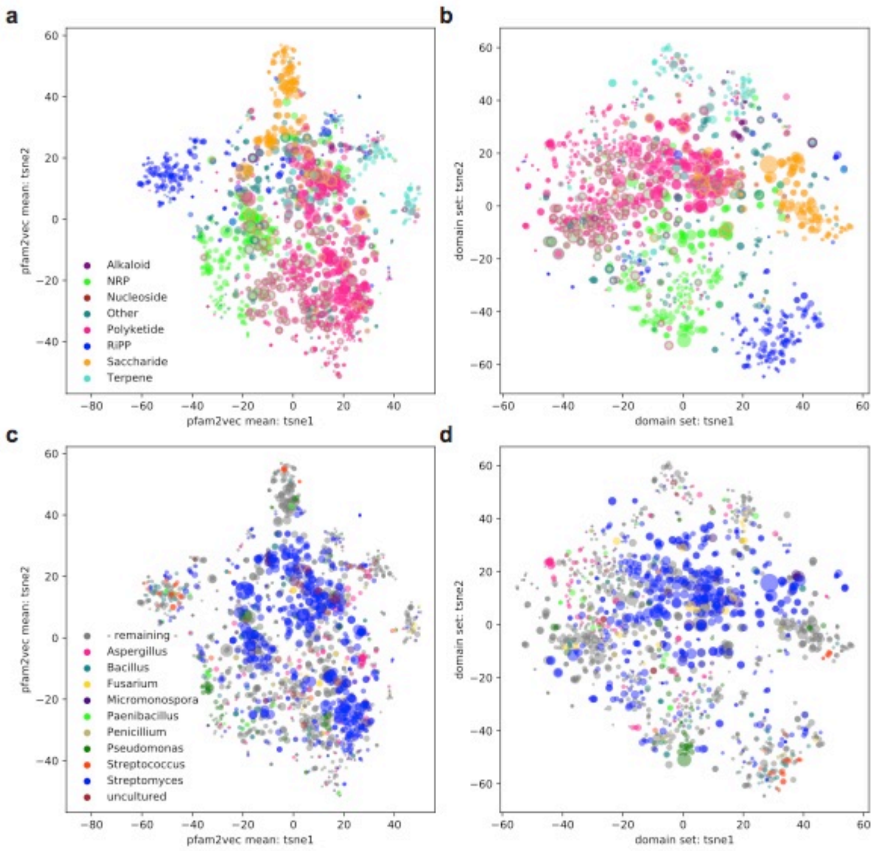


**Supplementary Figure S3.** t-Distributed Stochastic Neighbor Embedding (t-SNE) of all 1406 BGCs from the MIBiG database (Minimum Information about a Biosynthetic Gene cluster) using (**a**) average pfam2vec vector, colored by BGC class (**b**) one-hot-vector, colored by BGC class (**c**) same as '**a**', but colored by species (**d**) same as '**b**', but colored by species.

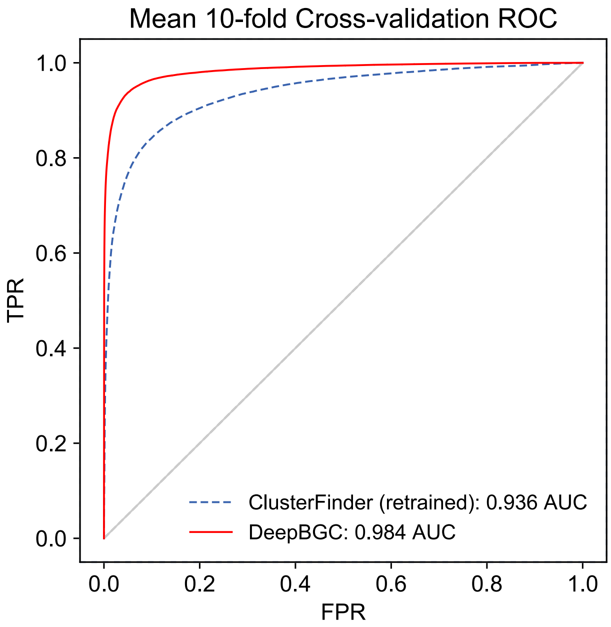


**Supplementary Figure S4.** Cross-validation ROC. Curves reflecting average performance of: (green) Retrained HMM model and (red) DeepBGC BiLSTM following 10-fold cross validation analysis. A total of 1406 BGCs from MiBIG database were used as a positive set alongside 10128 artificially created non-BGC negative set.

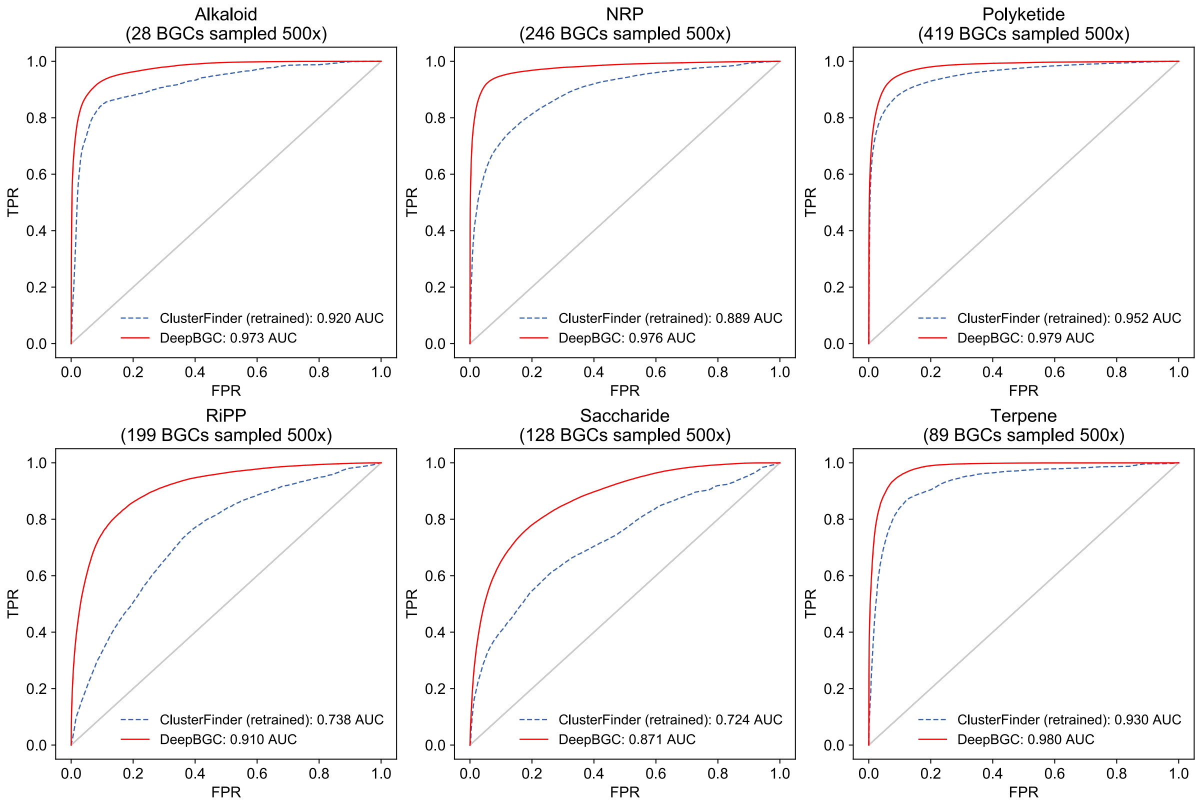


**Supplementary Figure S5.** ROC curves reflecting average performance of: (green) Retrained HMM model and (red) DeepBGC model following "Leave-Class-Out" analysis. Individual classes performance are shown here: Alkaloid, Non-Ribosomally synthesized Peptides (NRP), Polyketides (PKS), Ribosomally Synthesized and Post-translationally modified Peptides (RiPP) Saccharides and Terpenoids.

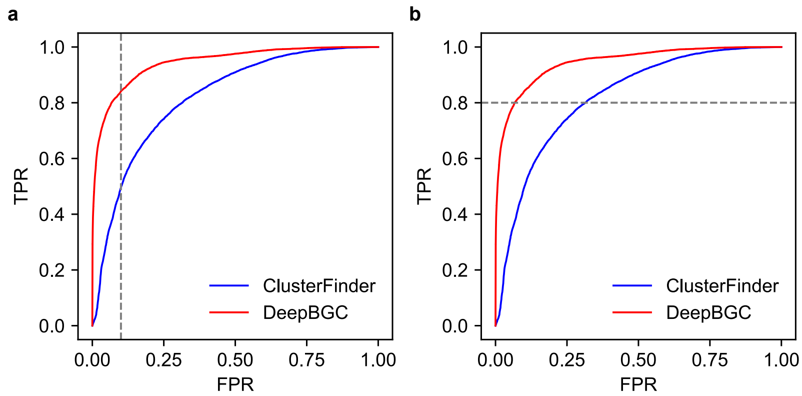


**Supplementary Figure S6.** ROC curves used to define (**a**) 10% FPR (**b**) 80% TPR domain level thresholds for DeepBGC (red) and ClusterFinder (blue) algorithms. The respective confusion matrix is available in Supplementary Table S7. For both models, the ROC was derived using 7 species from the test set of the first split in bootstrap validation.

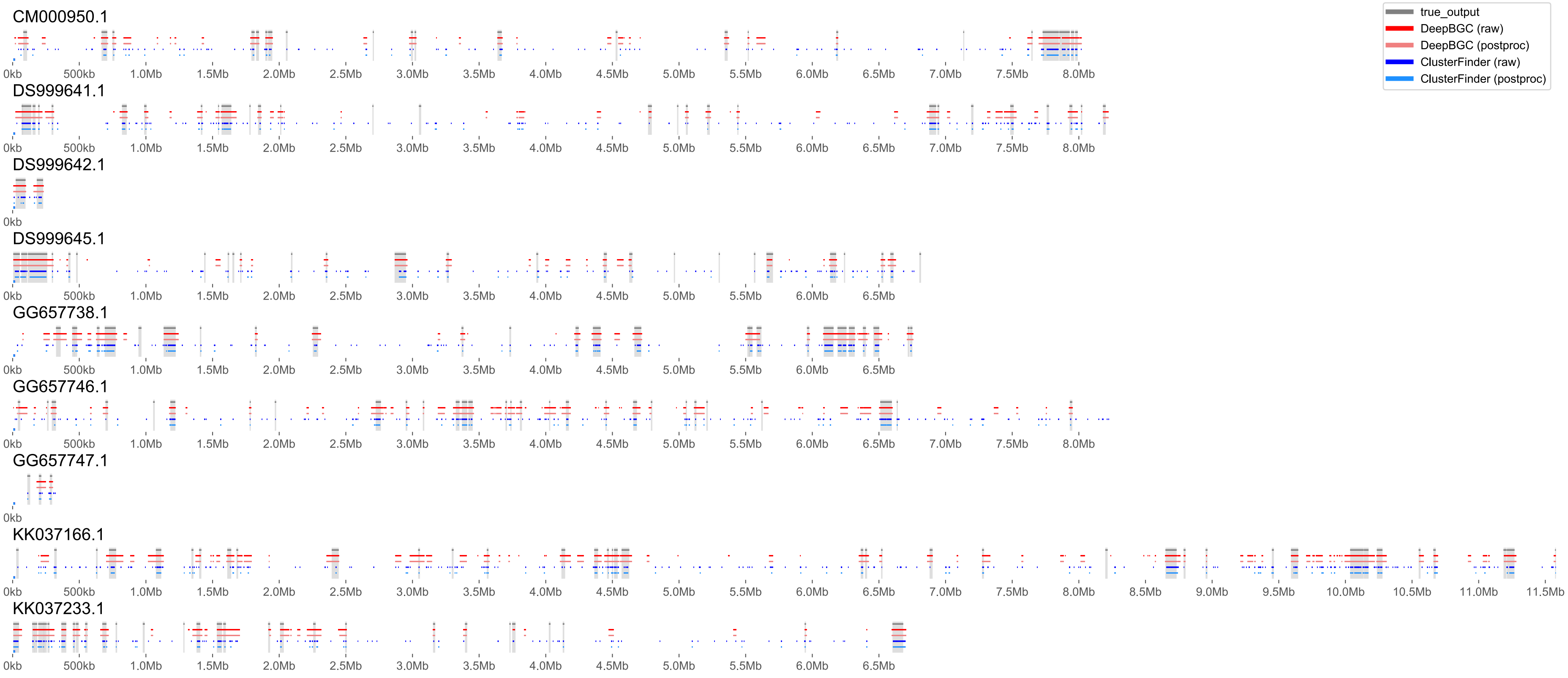


**Supplementary Figure S7.** A contig view (X-axis genomic coordinates) of true predictions (grey shade and bar), ClusterFinder raw and post-processed predictions (dark and light blue), DeepBGC raw and post-processed (dark and light red) for a total of 7 tested species present in 9 contigs. A 10% FPR threshold was applied to both models.

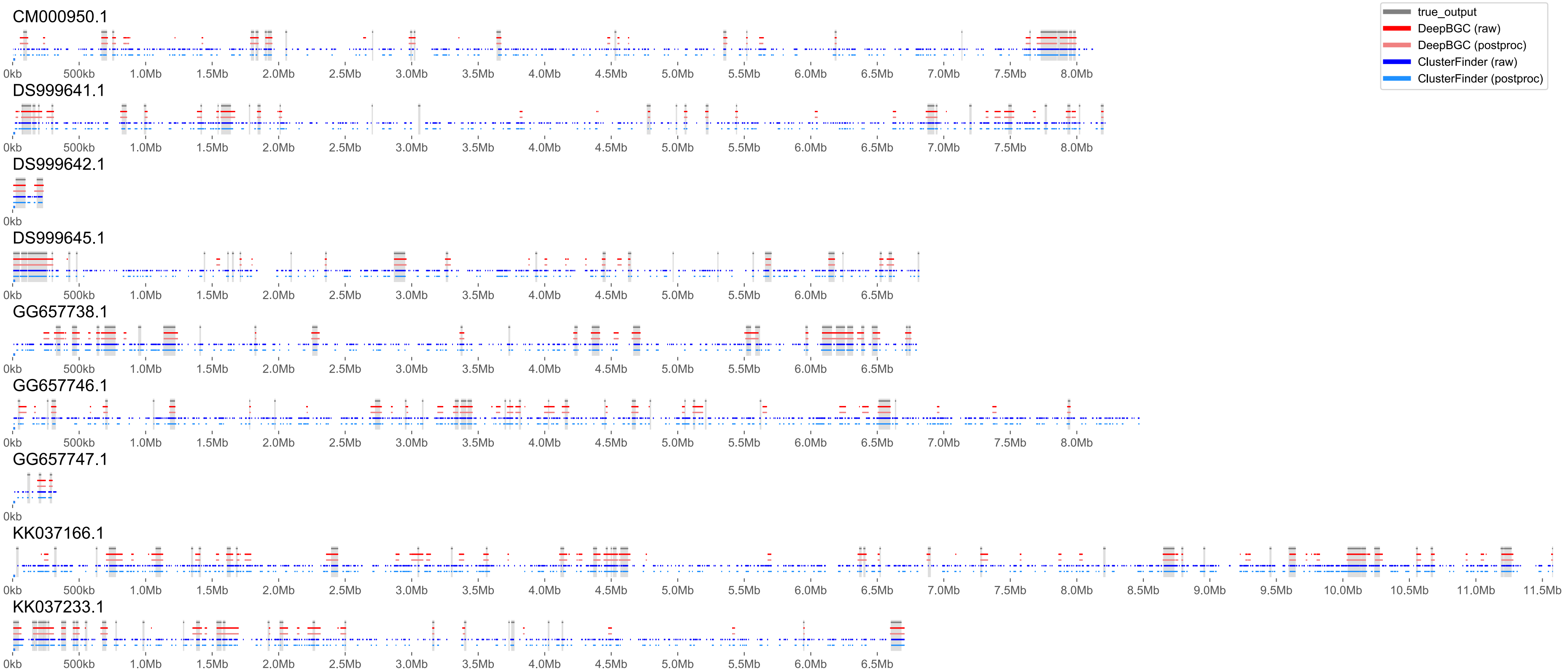


**Supplementary Figure S8.** A contig view (X-axis genomic coordinates) of true predictions (grey shade and bar), ClusterFinder raw and post-processed predictions (dark and light blue), DeepBGC raw and post-processed (dark and light red) for a total of 7 tested species present in 9 contigs. An 80% TPR threshold was applied to both models.

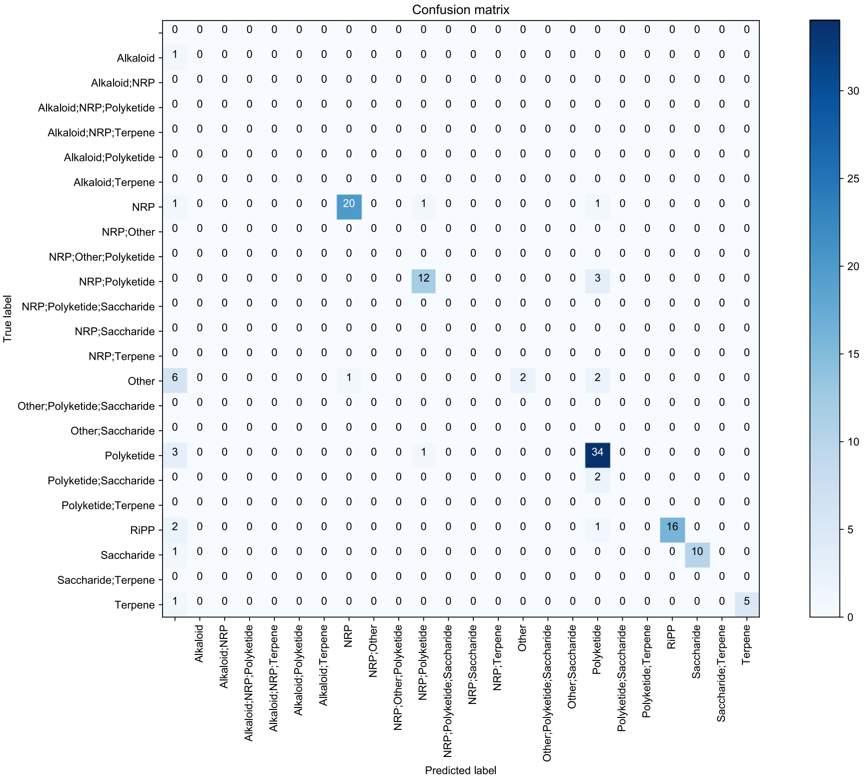


**Supplementary Figure S9.** Confusion matrix of random forest classifier predictions of BGC classes. The classifier was trained using 1355 MIBiG labeled BGCs belonging to one or more compound classes including Polyketides (PKS), Non-Ribosomally synthesized Peptides (NRP), Ribosomally Synthesized and Post-translationally modified Peptides (RiPP), Saccharides, Terpenes, Alkaloids, and those belonging to other rarer classes ("Other"). Respective Area Under the Curve (AUC) values are available in Table 1.

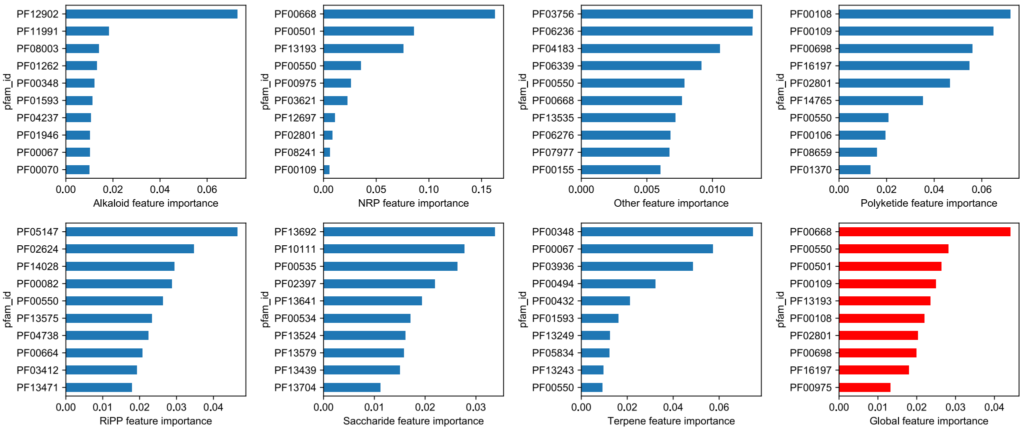


**Supplementary Figure S10.** Feature importance as extracted from random forest classifier for BGC classes. The classifier was trained using 1355 MIBiG labeled BGCs belonging to one or more compound classes including Polyketides (PKS), Non-Ribosomally synthesized Peptides (NRP), Ribosomally Synthesized and Post-translationally modified Peptides (RiPP), Saccharides, Terpenes, Alkaloids, and those belonging to other rarer classes ("Other"). The Feature importance score (X-axis) alongside Pfam IDs (Y-axis) are provided for each class. Global feature importance is also provided at the bottom right panel (red).

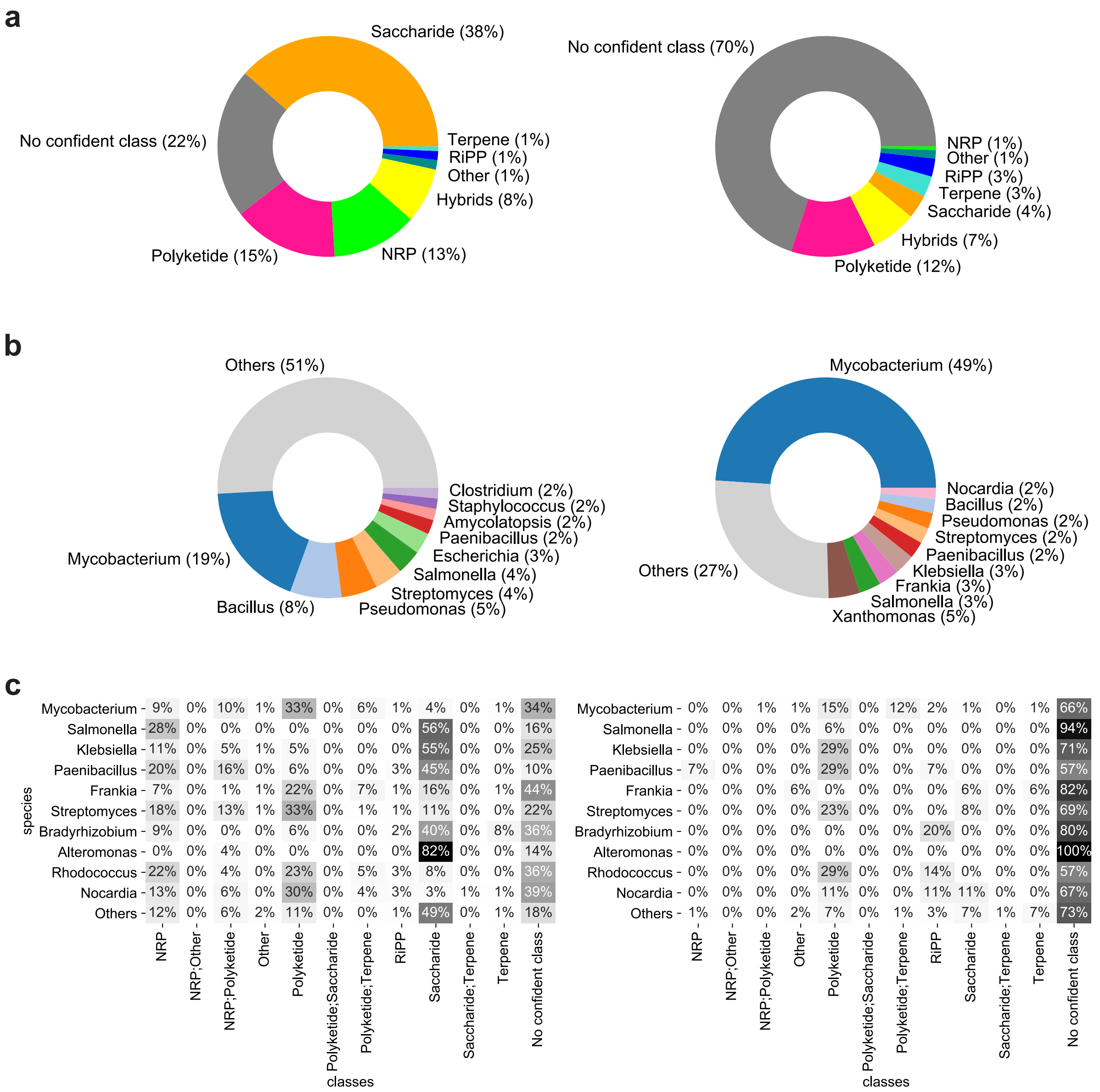


**Supplementary Figure S11.** Distribution of 10926 total BGCs predicted by DeepBGC (left) and 566 novel BGCs that could not be predicted by other models (right) according to their (**a**) class (BGCs with multiple classes predicted are grouped into "Hybrids") (**b**) underlying species (**c**) combination of class and underlying species. Only top 10 species are shown as ranked by number of BGCs predicted, remaining values are grouped into "Others".

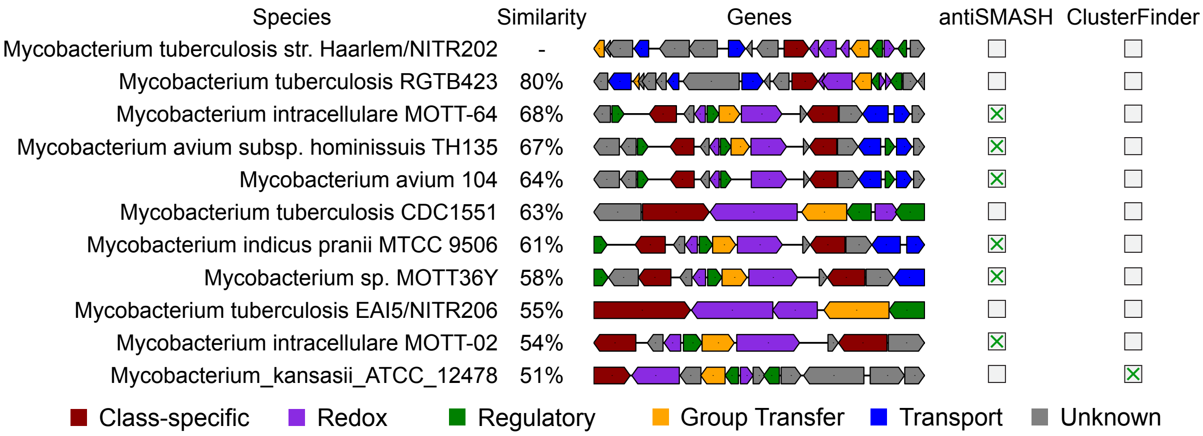


**Supplementary Figure S12.** Novel *Mycobacterium tuberculosis* DeepBGC candidate (first row) and top 10 most similar DeepBGC candidates (remaining rows). Candidates are found in 10 different strains of 5 different species of *Mycobacterium* (*tuberculosis*, *intracellulare*, *avium*, *indicus pranii* and *kansasii*, first column). Similarity was defined as cosine similarity of one-hot-vector representation (second column). Genes of each candidate are colored based on the underlying domain type (third column). Six of these candidates were detected also by antiSMASH (fourth column) and one was detected also by ClusterFinder (last column).
